# Access and local retention control neutrophil function in retinal regeneration

**DOI:** 10.64898/2026.11.16.766911

**Authors:** Yi Li, Tao Xu, Yikun Zhi, Ran Zhao, Qianyi Lu, Hui Xu

## Abstract

Neutrophils are among the first immune cells to reach injured tissues, yet their contributions to regeneration range from essential to dispensable or even inhibitory across injury contexts. Whether this variability reflects differences in tissue access, post-entry function, or persistence remains unresolved. Here we show that neutrophils are required for regenerative responses in needle-poked, but not NMDA-injured, zebrafish retinas. In stab injury, where barrier disruption permits neutrophil entry, pharmacological neutrophil ablation suppresses Müller glia proliferation, progenitor formation, and neuronal regeneration. In contrast, in the NMDA injury model, neutrophils show minimal infiltration into the retinal parenchyma, and their depletion has no significant effect. We further show that Cxcl12-Cxcr4b signaling retains neutrophils at the stab injury site, and that manipulating retention correspondingly alters regenerative outcomes. At the injury site, neutrophils modulate microglial reactivity and inflammatory responses and provide Hb-egf as an early regeneration-promoting factor. These findings identify a previously unrecognized requirement for neutrophils in retinal regeneration and reveal a two-tiered control of their function: access to the injured tissue, and local retention that enables regenerative signaling.

**Significance Statement:** Neutrophils contribute to regeneration in some injury contexts but not others, yet what determines this difference remains poorly understood. Here we show that in the injured zebrafish retina, neutrophils are required for regeneration after stab injury but not after NMDA injury, where they were largely excluded from the retinal parenchyma. Once access is permitted, Cxcl12-Cxcr4b signaling retains neutrophils at the injury site, where they modulate microglial reactivity and inflammatory responses and provide Hb-egf to support Müller glia reprogramming. These findings suggest that neutrophil contributions to retinal regeneration are governed not by cell identity alone, but by the coupling of tissue access with local retention that enables regenerative signaling.

## 1. INTRODUCTION

Adult neural stem cells (NSCs) hold great promise for curing nerve injury or diseases, as they self-renew and generate newborn neurons to restore nervous system function. NSCs reside in a specialized stem cell niche, constituted by supporting cells, immune cells, the vascular system, and other non-cellular factors(1, 2), which is important for NSC maintenance, quiescence, proliferation and differentiation(3). Immune cells are increasingly recognized as active components of this niche, yet how individual immune cell types contribute to NSC activation and subsequent neuronal regeneration remains poorly understood.

The zebrafish retina is a powerful model to study these mechanisms due to its robust regenerative capacity. Retinal regeneration in zebrafish is dependent on Müller glia (MG), a type of NSC-like macroglia that reprogram and divide asymmetrically to generate MG-derived progenitor cells (MGPCs) while maintaining the MG pool(4, 5). MGPCs resemble late-stage retinal progenitors during development(6) and further proliferate and differentiate into retinal neurons to restore visual function. Although MG in the mammalian retina share many characteristics with those found in zebrafish, they possess limited regenerative capacity under normal conditions(7). Recent progress has been made in reprogramming mammalian MG into retinal neurons *in vivo*(8, 9), but human application remain hindered by technological immaturity and ethical concerns. A thorough understanding of the mechanisms governing MG reprogramming and neuronal regeneration in zebrafish is therefore important for developing new strategies for human retinal regeneration.

MG reprogramming in the zebrafish retina is regulated by a complex network of intrinsic factors and extracellular cues(4, 10). Among the extracellular cues, retinal inflammation and the innate immune responses play important roles in MG reprogramming and retinal regeneration(11–13). Following retinal injury, innate immune cells such as microglia and macrophage, become reactive and are recruited to the injury site within hours(14, 15). Activated microglia/macrophage clear dead cells and debris by phagocytosis and express various soluble factors to modulate the inflammatory and regenerative responses in the zebrafish retina (16–19). Unlike in zebrafish, the role of microglia in mammalian MG reprogramming appears controversial: one study reports that they suppress this process(20), while two other research suggest otherwise(21) (22), possibly reflecting differences in reprogramming strategy or experimental model. These discrepancies raise a broader question: what determines whether a given myeloid population contributes to MG reprogramming—its presence at the injury site, its functional state once there, or both?

Neutrophils, another innate immune cell type, are the first responders to tissue injury. Their recruitment to the injured retina is itself injury model-dependent: neutrophils infiltrate the retina after puncture injury but show no reactivity to sterile rod cell loss in zebrafish larvae(15). Our previous work also revealed that large numbers of neutrophils enter the stab-injured adult zebrafish retina and accumulate at the injury site within 3 hours post injury (hpi) (23). In other regenerating zebrafish tissues, neutrophils are required for spinal cord and heart regeneration, although their pro-regenerative function depends on timely resolution at the injury site(24, 25). However, whether neutrophils causally contribute to MG reprogramming and retinal regeneration rather than merely accompanying injury-induced inflammation, and whether tissue access and local retention govern this contribution remain to be directly tested. Here we investigate the neutrophil response and its role in MG reprogramming and retinal regeneration using two injury paradigms in adult zebrafish: needle-poke injury, which disrupts the blood-retinal-barrier (BRB) and permits neutrophil infiltration into the retina, and NMDA injury, in which neutrophils are largely excluded from the retinal parenchyma. Our findings demonstrate that neutrophils play an injury paradigm-dependent role in zebrafish retinal regeneration and suggest that modulating neutrophil access or retention may offer therapeutic opportunities for certain types of retinal injury.

## 2. MATERIALS AND METHODS

### 2.1 Animals and Eye Injury

Zebrafish used in this study were treated following the Guidelines for Animal Use and animal care of Nantong University. Adult Wild-type (AB) and *Tg(mpx:GFP)*(*26*) transgenic zebrafish (6 to 8 months old) were maintained in an automatic breeding system at 28 °C on 14 hr light: 10 hr dark cycles. The method of needle-poke injury of zebrafish retina has been described in our previous study(12). Briefly, zebrafish were anesthetized in 0.02% of Tricaine methane sulfonate (Cat # E20521, Sigma-Aldrich) in system water. Fish were then transfer to a wet sponge and under a dissecting microscope, the right eye were gently rotated in the socket to expose the back, and were stabbed four times (once in each quadrant) with a sterile 30 G needle. The needle was inserted through the sclera to the length of the bevel. Care was taken not to damage the optic nerve and vessels. The uninjured left eye was used as the negative control.

The retinal NMDA-injury was performed as described previously(23). Briefly, fish were anesthetized and 1 μl of 50 mM NMDA (N-methyl-D-aspartate, Cat # M3262, Sigma-Aldrich) in PBS were injected intravitreally through the front of the eye. Care was taken not to damage the retina or lens.

### 2.2 Drug treatment

To deplete neutrophils, zebrafish received daily intraperitoneal injection of 20 μl of AG-845 (Cat # HY-15844, MedChemExpress) or CP-724714 (Cat # HY-14674, MedChemExpress) for three days before retinal injury. Both reagents were dissolved in DMSO as 20 mM stock solutions. The working concentrations of AG-845 and CP-724714 are 0.5 mM and 1 mM, respectively. AMD3100 (Cat # HY-10046, MedChemExpress) was dissolved as 90 mM stock solution in ethanol and fish received daily intravitreal injection of 1 μl of 1 mM AMD3100 until harvest.

Recombinant zebrafish Cxcl12a (SDF-1α) protein (Cat# abx178492, Abbexa) was dissolved as 1 mg/ml stock solution in PBS. 1 μl of 5 ng/μl Cxcl12a was injected intravitreally through the front of the eye at the time of retinal injury. Recombinant human HB-EGF protein (Cat# HY-P7400, MedChemExpress) was dissolved as 0.2 mg/ml stock solution in PBS. Zebrafish received intravitreal injection of 1 μl of 200 ng/μl of HB-EGF as described previously(27). To enhance the immune response in the retina, fish received intravitreal injection of 1 μl of 10 mg/ml of Zymosan A (Cat# HY-W250113, MedChemExpress) at the time of retinal injury.

### 2.3 Single-cell RNA sequencing (scRNA-seq) and analysis

Single cell suspensions were prepared from uninjured (0 d) and stab-injured (1 d, 2 d, and 4 d) adult zebrafish retinas. Detailed methods of sample preparation, library construction, sequencing and analysis have been described previously(18). The raw scRNA-seq data was deposited in the Genome Sequence Archive at the National Genomics Data Center, China National Center for Bioinformation, under accession number CRA045511 (https://ngdc.cncb.ac.cn/gsa/).

### 2.4 Immunofluorescence, Lineage Tracing, and TUNEL staining

Methods for retinal tissue preparation, cryosection, and immunofluorescence staining were described previously(12). The following primary antibodies were used: rabbit anti-GNAT2 (1:200, Cat # PM075, MBL Life Science); mouse anti-HuC/D (1:500, Cat # A-21271, Thermo Fisher Scientific).

1 mg of DyLight 594-labeled isolectin B4 (IB4; Cat # DL-1207-.5, Vector laboratories) was intravitreally injected one day before sacrifice to label microglia in the zebrafish retina. To label proliferating cells in the retina, 5-Ethynyl-2’-deoxyuridine (EdU) staining was performed using a BeyoClickTM EdU-488 Cell Proliferation Detection Kit (Cat# C0071L, Beyotime, Shanghai, China). For MGPC lineage tracing, fish received intraperitoneal injection of 20 mM EdU solution (20 μl per fish, Cat# HY-118411, MedChemExpress) at 4 dpi. The TUNEL staining was performed using the One Step TUNEL Apoptosis Assay Kit (Red Fluorescence) (Cat# C1089, Beyotime) according to manufacturer’s protocol.

### 2.5 Reverse transcription PCR (RT-PCR) and Quantitative PCR (qPCR)

Fish retinas were collected and total RNA was isolated using the RNA isolater Total RNA Extraction Reagent (Cat # R401-01, Vazyme, Nanjing, China). RT-PCR was performed as described previously(28). qPCR was performed in triplicate using methods described previously(29) and the biological replicate *n* ≥ 3. Relative mRNA levels were calculated using the ΔΔCt method(30) and normalized to the level of a house-keeping gene *rpl13*. Primers used in RT-PCR and qPCR were listed in **Table 1**.

**Table 1.**
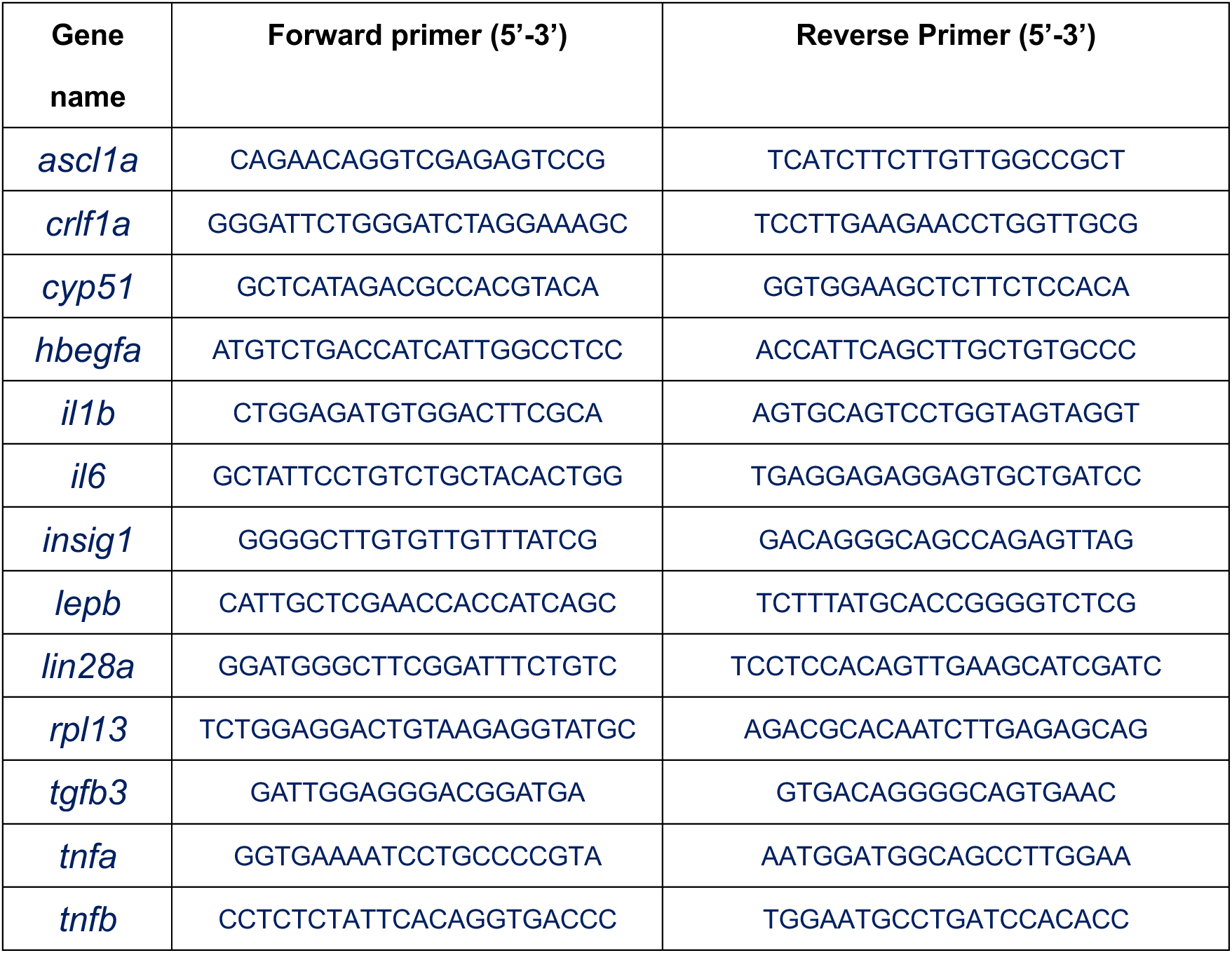
Primers used for RT-PCR and qPCR.

### 2.6 Imaging and cell counting

Images of retinal cryosections were acquired using a Zeiss Imager M2 fluorescence upright microscope (Carl Zeiss AG) with 5x,10x or 20x objectives. Cell counting of the retina was performed using the Cell Counter plugin in the ImageJ Software (https://imagej.net/ij/).

### 2.7 Statistics

All experimental data were analyzed using the GraphPad Prism 11.0 software. A student’s *t-test* (2 tails, unpaired) and one-way analysis of variance (ANOVA) followed by a Dunnett’s multiple comparison test were used for the statistical analyses. The difference was considered significant at \**p* < 0.05, \*\**p* < 0.01; \*\*\**p* < 0.001. Representative results are shown as mean±SEM.

## 3. RESULTS

### 3.1 Neutrophil dynamics in the stab-injured zebrafish retina

To investigate neutrophil dynamics in the retina following stab-injury, we took advantage of the *Tg(mpx:GFP)* transgenic zebrafish line which specifically labels neutrophils with GFP, and examined neutrophil together with microglia in the injured retina at 1, 2, 3, 6, 12 and 24 hours post injury (hpi). In the uninjured intact retina, neutrophils were absent (Figure 1A, 0 h). After needle-poked injury, neutrophils entered the retina as early as 1 hpi, peaked at 3 hpi, and gradually decreased from 6 to 24 hpi (Figure 1A,B). In contrast, microglial accumulation at the injury site, identified by IB4 staining, was not evident until 3 hpi and further increased from 6 to 24 hpi (Figure 1A,B). By 24 hpi, the majority of innate immune cells at the injury site were microglia. These results indicate an early but transient neutrophil response in blood-retina-barrier (BRB)-disrupted zebrafish retinas.

**Figure 1.**
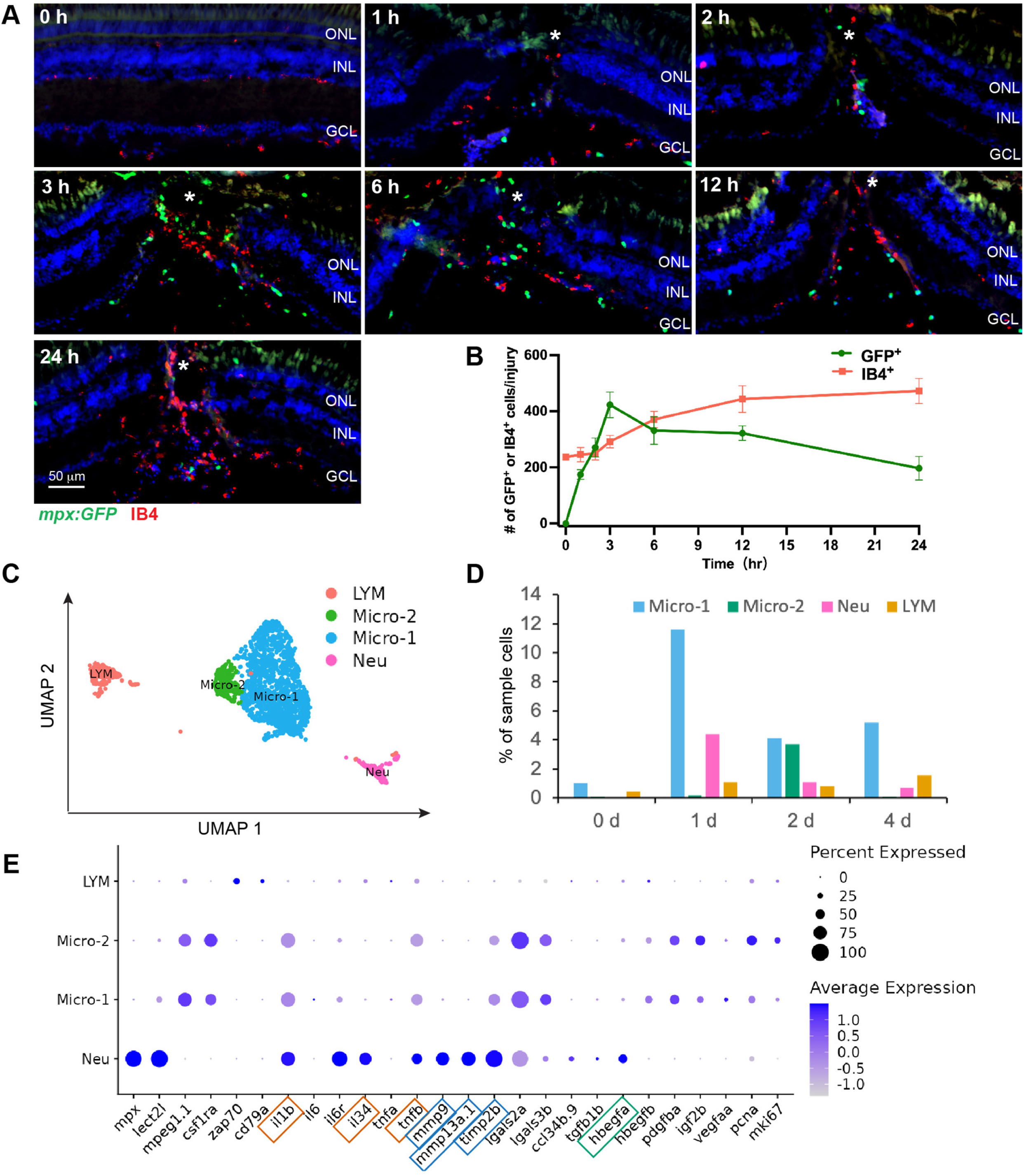
Neutrophil dynamics and molecular signature in stab-injured zebrafish retinas. (A) Fluorescence microscopy showing the presence of neutrophils (GFP^+^, green) and microglia (IB4^+^, red) in uninjured (0 h) or stab-injured *Tg(mpx:GFP)* zebrafish retinas at indicated timepoints. (B) Quantification of neutrophil and microglia numbers in (A). (C,D) scRNA-seq analysis showing the cell clustering and composition of retinal immune cells in uninjured (0 d) and stab-injured (1 d, 2 d, 4 d) zebrafish retinas. (E) Dotplot showing the molecular signature of retinal immune cells. White asterisks, site of the needle-poke injury. ONL, outer nuclear layer; INL, inner nuclear layer; GCL, ganglion cell layer; LYM, lymphocyte; Micro, microglia; Neu, neutrophil.

We next examined the composition and molecular signature of retinal immune cells in intact and stab-injured zebrafish retinas by analyzing previously performed single-cell RNA sequencing (scRNA-seq) data from our lab (Li et al, bioRxiv 2026.06.10.731489). Using known cell type markers, we identified neutrophils, two microglia subtypes (micro-1 and micro-2), and lymphocytes in the retina (Figure 1C-E). Consistent with these findings, neutrophils were not detected in the uninjured retina (Figure 1D, 0 d), and their numbers decreased substantially after 1 day post injury (dpi) (Figure 1 D). Analysis of gene expression within the immune populations showed that neutrophils expressed higher levels of inflammatory cytokines (*il1b*, *tnfb*) and matrix metalloproteinases (*mmp9* and *mmp13a.1*) than microglia and lymphocytes (Figure 1E, red and blue boxes), suggesting that they act as key effector cells driving acute retinal inflammation and matrix remodeling. Unlike microglia which expressed several growth factors (*hbegfb*, *pdgfba*, *igf2b* and *vegfaa*), only one growth factor—*hbegfa,* was identified expressed by neutrophils in the retina (Figure 1E, green box).

### 3.2 Neutrophils are required for the regenerative responses in stab-injured retinas

To investigate the potential role of neutrophils in zebrafish retinal regeneration, we pharmacologically ablated neutrophils by treating the fish with Erbb2 inhibitors AG-845 or CP-724714 (hereafter AG or CP; experimental timeline in Figure 2A), both of which has been shown to accelerate neutrophil apoptosis and promote neutrophilic resolution *in vitro* and *in vivo* (31). Both AG and CP treatment significantly reduced the number of neutrophils at the injury site of needle-poked retina (Figure 2A,B), confirming their effectiveness in depleting neutrophil in adult zebrafish. Neither AG nor CP treatment caused a significant increase in the number of TUNEL^+^ cells in either the injured or uninjured retinal region (Figure 2C), suggesting that neither treatment has obvious cytotoxicity to retinal cells.

**Figure 2.**
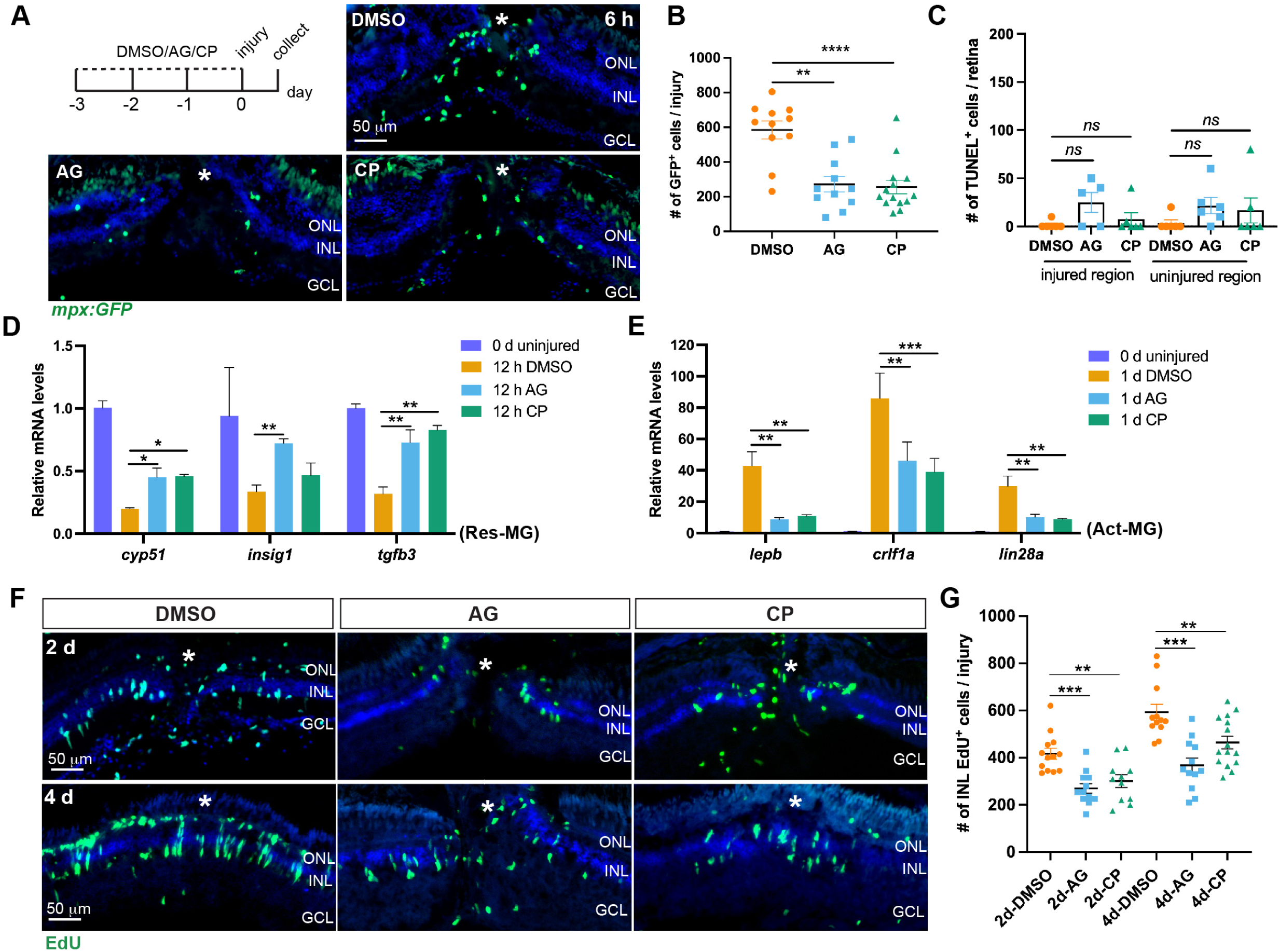
Neutrophils are required for the regenerative response in stab-injured zebrafish retina. (A) Experimental timeline and fluorescence microscopy showing the accumulation of neutrophils (green) at the injury site in stab-injured *Tg(mpx:GFP)* zebrafish retinas at 6 hpi. White *, site of the needle-poke injury. (B) Quantification of neutrophil numbers at the injury site in (A). (C) Quantification of the TUNEL^+^ cells in DMSO or AG/CP-treated retinas at 6 hpi. (D,E) qPCR analysis of the retinal expression of marker genes of Res-MG (D) and Act-MG (E). *n* ≥ 3 for each group. (F) EdU immunofluorescence showing the presence of proliferating cells in DMSO- or AG/CP-treated retinas at 2 and 4 dpi. (G) Quantification of the number of INL EdU^+^ cells in (F). White asterisks, site of the needle-poke injury. ONL, outer nuclear layer; INL, inner nuclear layer; GCL, ganglion cell layer.

Because neutrophil infiltration into the injured retina is an early event that overlaps with the period of MG reprogramming (∼0-1 dpi), we first examined the effects of neutrophil ablation on MG status transition by measuring the expression levels of MG status markers (18). qPCR showed that after retinal injury, expression of resting-MG markers such as *cyp51*, *insig1* and *tgfb3* was largely downregulated, whereas AG or CP treatment significantly attenuated this downregulation (Figure 2D). Conversely, both AG and CP treatment significantly suppressed the induction of activated-MG markers *lepb*, *crlf1a*, and *lin28a* in the retina (Figure 2E). These results indicate that neutrophils are indispensable for MG reprogramming in stab-injured zebrafish retina.

We next examined the effect of neutrophil ablation on MG proliferation and subsequent MGPC formation. In stab-injured zebrafish retinas, MG enter the cell cycle at about 2 dpi and can be identified by BrdU or EdU labeling in the inner nuclear layer (INL) (17, 32). MG-derived cells further proliferate into neurogenic MGPC clusters, which can be identified in the INL by the same cell cycle markers at 4 dpi (12, 17, 32). Immunofluorescence staining showed that AG or CP treatment significantly decreased the number of EdU^+^ cells in the INL at 2 dpi (Figure 2F upper panel, Figure 2G), suggesting reduced MG proliferation. Subsequently, AG or CP treatment led to a significantly reduced number of MGPCs at 4 dpi (Figure 2F lower panel, Figure 2G). Taken together, these results indicate that neutrophils are required for the regenerative responses in stab-injured zebrafish retinas.

### 3.3 Neutrophil depletion impairs the regeneration of retinal neurons

To investigate how neutrophil depletion affects the regeneration of retinal neurons, control or AG/CP-treated fish received an intraperitoneal EdU injection at 4 dpi to label proliferating MGPCs, and their progeny cells were analyzed at 7 and 30 dpi (Figure 3A, experimental timeline). EdU immunofluorescence at 7 dpi showed that the number of MGPC progeny cells in AG- or CP-treated retinas were lower than that of control (Figure 3 A,B). Furthermore, the percentage of early differentiating MGPCs, identified by EdU and HuC/D double staining at this timepoint(12, 33), was also significantly lower in AG- or CP-treated retinas than in controls (Figure 3A,C). These data suggest that neutrophil depletion not only reduced the overall MGPC population but may also have negatively affected their early differentiation. At 30 dpi, AG- or CP-treated retinas had fewer MGPC progeny cells across all three nuclear layers than controls, as shown by the EdU staining (Figure 3D,E). Co-staining with cell type markers demonstrated a decreased number of regenerated neurons, including photoreceptors (Gnat2^+^ EdU^+^), amacrine cells (HuC/D^+^ EdU^+^ in the INL), and retinal ganglion cells (RGCs, HuC/D^+^ EdU^+^ in the GCL) in AG- or CP-treated retinas at 30 dpi (Figure 3D,F). Together, these data indicate that neutrophil depletion negatively affects the regeneration of retinal neurons in stab-injured zebrafish retinas.

**Figure 3.**
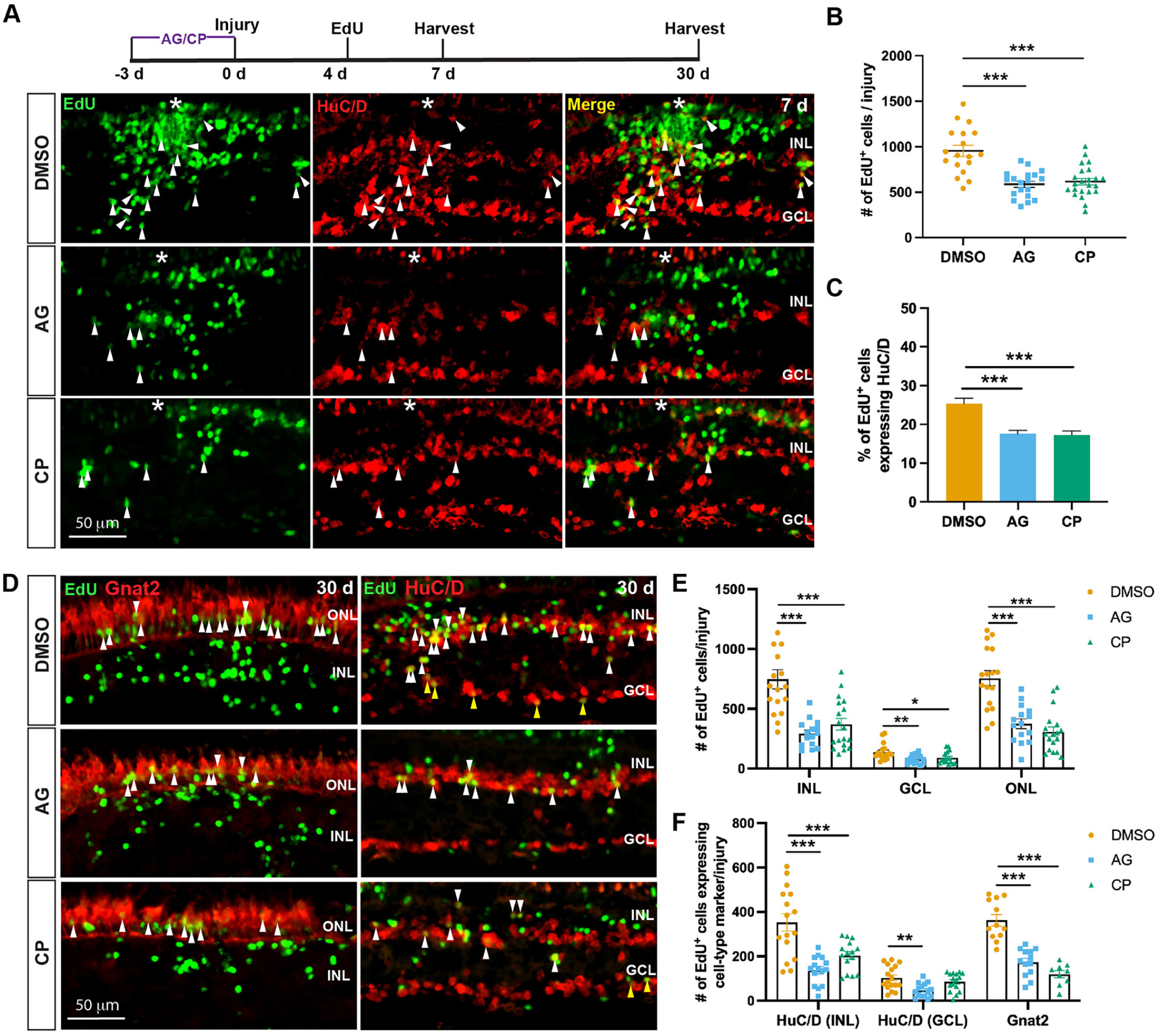
Neutrophil depletion results in impaired retinal neuron regeneration in stab-injured retinas. (A) Experimental timeline of the lineage-tracing experiments, and fluorescence microscopy showing the early differentiation of MGPCs in DMSO or AG/CP-treated retinas at 7 dpi. Arrowheads, EdU^+^ HuC/D^+^ cells. (B,C) Quantification of the number of EdU^+^ cells per injury and the proportion of EdU^+^ cells expressing HuC/D in (A). (D) Co-staining of cell type markers with EdU to identify MGPC-derived cones (Gnat2^+^ EdU^+^), amacrine cells (HuC/D^+^ EdU^+^ in the INL) and retinal ganglion cells (RGCs, HuC/D^+^ EdU^+^ in the GCL) at 30 dpi in stab-injured retinas. (E,F) Quantification of the total number of MGPC progenies and regenerated retinal neurons at 30 dpi in (D). White asterisks, site of the needle-poke injury. ONL, outer nuclear layer; INL, inner nuclear layer; GCL, ganglion cell layer.

### 3.4 Neutrophil depletion has no significant effect on the regenerative response in NMDA-injured retinas

A previous study reported that neutrophils respond to retinal puncture wounds that break the BRB but not to laser-induced rod cell loss in which the BRB remains intact in zebrafish larvae(15). This raised an interesting question of whether neutrophils regulate the regenerative response in an injury paradigm-dependent manner. To address this question, we examined the neutrophil reactivity and retinal regenerative response in NMDA-injured retina, a BRB intact model. In uninjured control retinas, almost no neutrophils were visible in the retina or vitreous (Figure 4A), whereas some were present in the iris (Figure 4A boxed area; Figure 4A’). Following NMDA injury, no obvious neutrophil infiltration into the retina was observed, and few neutrophils were present in the injured retinas (Figure 4A-E). In contrast, neutrophil numbers in the vitreous and iris increased from 3 to 12 h (Figure 4A-E, A’-D’), likely as a result of NMDA injection through the front of the eye. These results suggest that neutrophils are largely excluded from the injured retinal parenchyma after NMDA injury.

**Figure 4.**
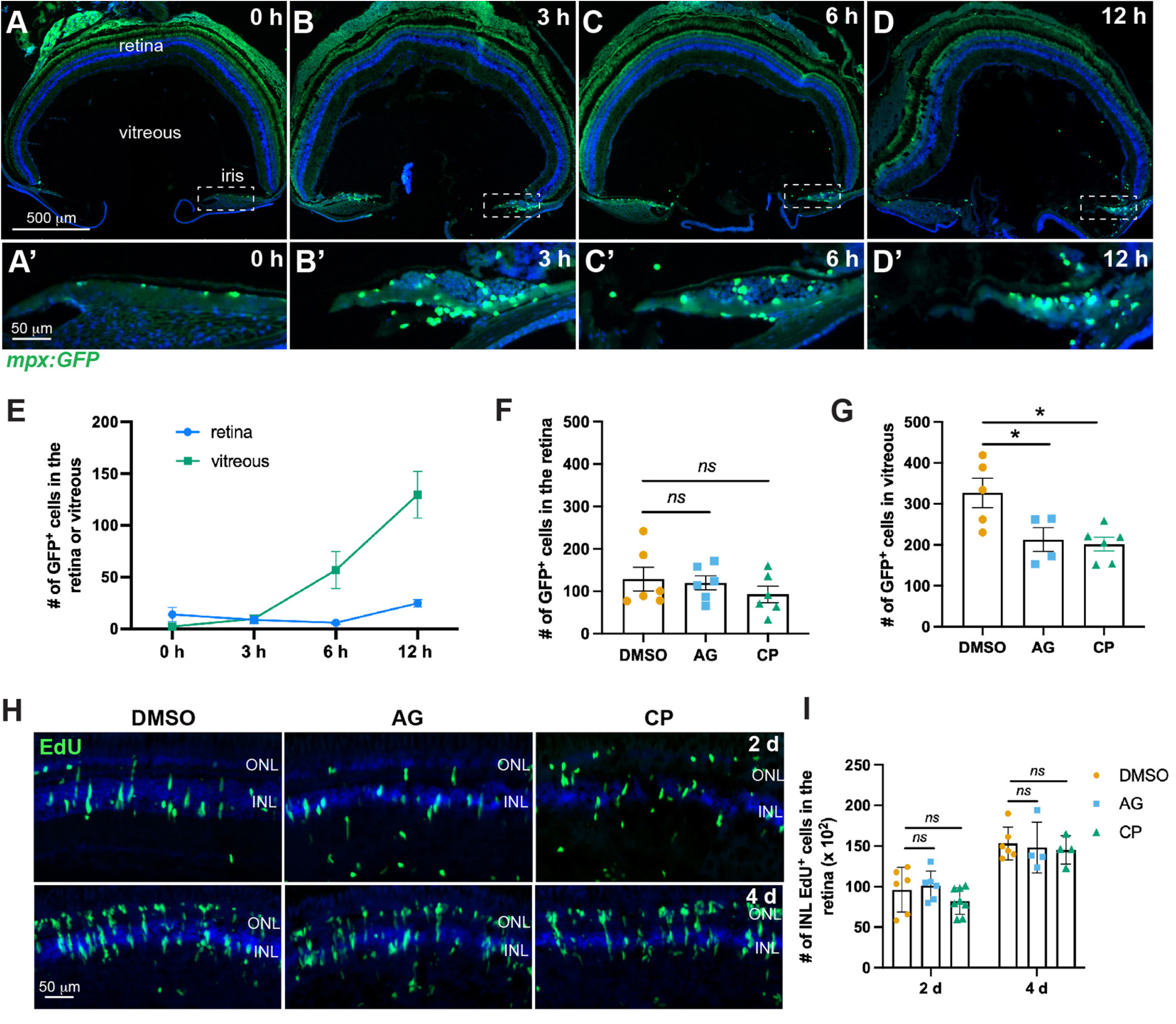
Neutrophils are not required for the regenerative response in NMDA-injured retinas. (A-D) Fluorescence microscopy showing the presence of neutrophils in uninjured or NMDA-injured *Tg(mpx:GFP)* zebrafish eyes at indicated timepoints. Some neutrophils can be seen in the iris (dashed box). (A’-D’) Zoomed in images of the iris in (A-D). (E) Quantification of the neutrophil number in the retina and vitreous. (F,G) Quantification of the neutrophil number in DMSO- or AG/CP-treated retina and vitreous at 6 hpi. (H) EdU immunofluorescence showing the presence of proliferating MG (2 dpi) and MGPCs (4 dpi) in DMSO- or AG/CP-treated retinas. (I) Quantification of the number of INL proliferating cells per retina in (H). ONL, outer nuclear layer; INL, inner nuclear layer.

We next investigated the effect of AG or CP treatment on neutrophil number and retinal regenerative responses in this model. Although intraperitoneal injection increased basal neutrophil numbers in all three groups (DMSO, AG, and CP), AG or CP treatment did not reduce neutrophil numbers in the retina compared to the DMSO control at 6 hpi (Figure 4F). In contrast, AG or CP treatment significantly decreased the number of neutrophils in the vitreous at this time point (Figure 4G). EdU immunofluorescence showed that AG or CP treatment had no significant effect on MG proliferation or MGPC formation at 2 and 4 dpi, respectively (Figure 4H,I). This result is consistent with the minimal neutrophil infiltration into the retinal parenchyma after NMDA injury and suggests that tissue access is a key determinant of the role of neutrophils in retinal regeneration.

### 3.5 Manipulating neutrophil retention in stab-injured retinas affects the regenerative responses

The Cxcl12-Cxcr4 chemokine signaling is known to retain neutrophils in the bone marrow and at inflammatory sites(34, 35), and our recent scRNA-seq study showed that neutrophils in stab-injured zebrafish retina express high levels of Cxcr4b (Li et al, bioRxiv 2026.06.10.731489). We therefore ask whether manipulating neutrophil retention in the injured retina would affect the regenerative responses. For this purpose, fish received intravitreal injection of a selective Cxcr4 antagonist AMD3100(36), or a recombinant zebrafish Cxcl12a protein, to suppress or enhance this chemokine signaling, respectively. AMD3100-treatment had no effect on initial neutrophil infiltration into the retina at 1 and 2 hpi but significantly reduced neutrophil numbers from 3 to 24 hpi compared to the ethanol control (Figure 5A), indicating an enhanced neutrophil resolution. Conversely, Cxcl12a injection did not affect early neutrophil infiltration into the retina at 2 and 3 hpi but significantly increased neutrophil numbers during the resolution phase from 6 to 24 hpi (Figure 5B). These results are consistent with a neutrophil retention effect of the Cxcl12-Cxcr4 signaling in stab-injured retinas.

**Figure 5.**
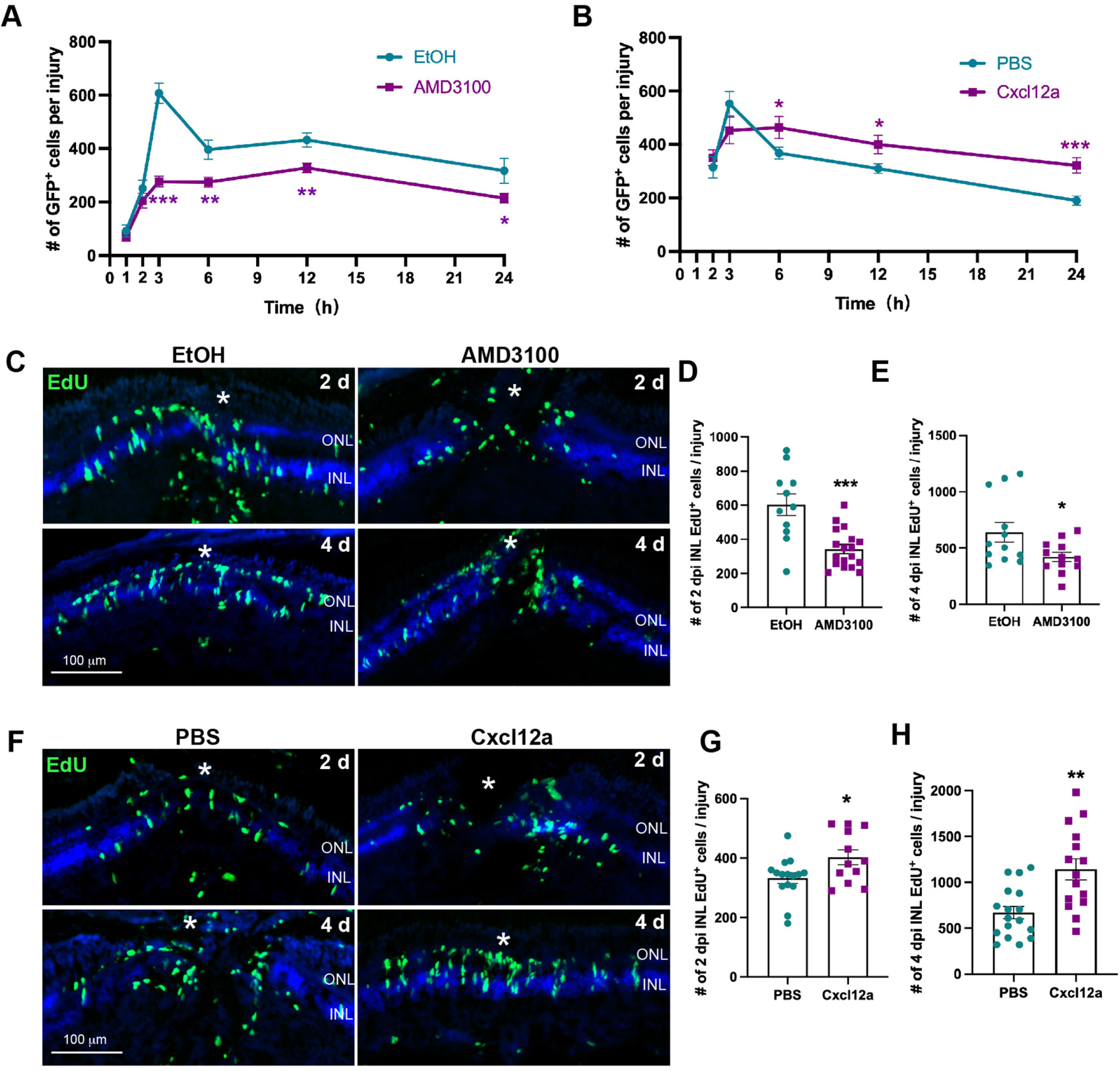
Neutrophilic retention by the Cxcl12-Cxcr4 signaling promotes the retinal regenerative responses in stab-injured retina. (A,B) Quantification of the neutrophil number per injury in control, AMD3100- or Cxcl12a-treated retinas at indicated timepoints. (C) EdU immunofluorescence showing the presence of proliferating MG (2 dpi) and MGPCs (4 dpi) in control (EtOH) or AMD3100-treated retinas. (D,E) Quantification of the number of INL proliferating cells per injury in (C). (F) EdU immunofluorescence showing the presence of proliferating MG (2 dpi) and MGPCs (4 dpi) in control (PBS) or Cxcl12a-treated retinas. (G,H) Quantification of the number of INL proliferating cells per injury in (F). White asterisks, site of the needle-poke injury. ONL, outer nuclear layer; INL, inner nuclear layer.

We next examined the effect of manipulating neutrophil retention on the regenerative responses in stab-injured retinas. EdU immunofluorescence staining showed that reducing neutrophil retention with AMD3100 significantly decreased the number of proliferating MG at 2 dpi and the number of MGPCs at 4 dpi (Figure 5C-E). Conversely, enhancing neutrophil retention by intravitreal Cxcl12a injection led to increased MG proliferation and subsequent MGPC formation compared to PBS controls (Figure 5F-H). Taken together, these results demonstrate that the Cxcl12-Cxcr4 signaling retains neutrophils in the injured retina and thereby enhances the regenerative responses.

### 3.6 Neutrophils regulate microglial reactivity and the inflammatory response in stab-injured retinas

Because neutrophils and microglia both accumulate at the injury site during the early phase after stab-injury, we hypothesized that neutrophils may regulate microglial reactivity and the retinal inflammatory response, both of which are essential for MG reprogramming and proliferation. Indeed, IB4 immunofluorescence showed that the number of microglia accumulated at the injury site in AG- or CP-treated retinas was significantly lower than that of control at 1 and 2 dpi (Figure 6A,B), suggesting reduced microglial reactivity in neutrophil-depleted retinas. qPCR revealed that AG or CP treatment reduced the expression levels of typical inflammatory cytokines such as *il1b*, *il6*, *tnfa* and *tnfb* in the injured retina (Figure 6C), indicating a dampened inflammatory response. To investigate whether neutrophil regulate the retinal regenerative response through modulation of inflammation, neutrophil-depleted retinas were treated with zymosan A to enhance the immune response. EdU immunofluorescence showed that zymosan A injection restored defective MGPC formation in AG- or CP-treated retinas (Figure 6D,E). These results suggest that neutrophils regulate the regenerative response by modulating retinal microglial reactivity and the inflammatory response in stab-injured retinas.

**Figure 6.**
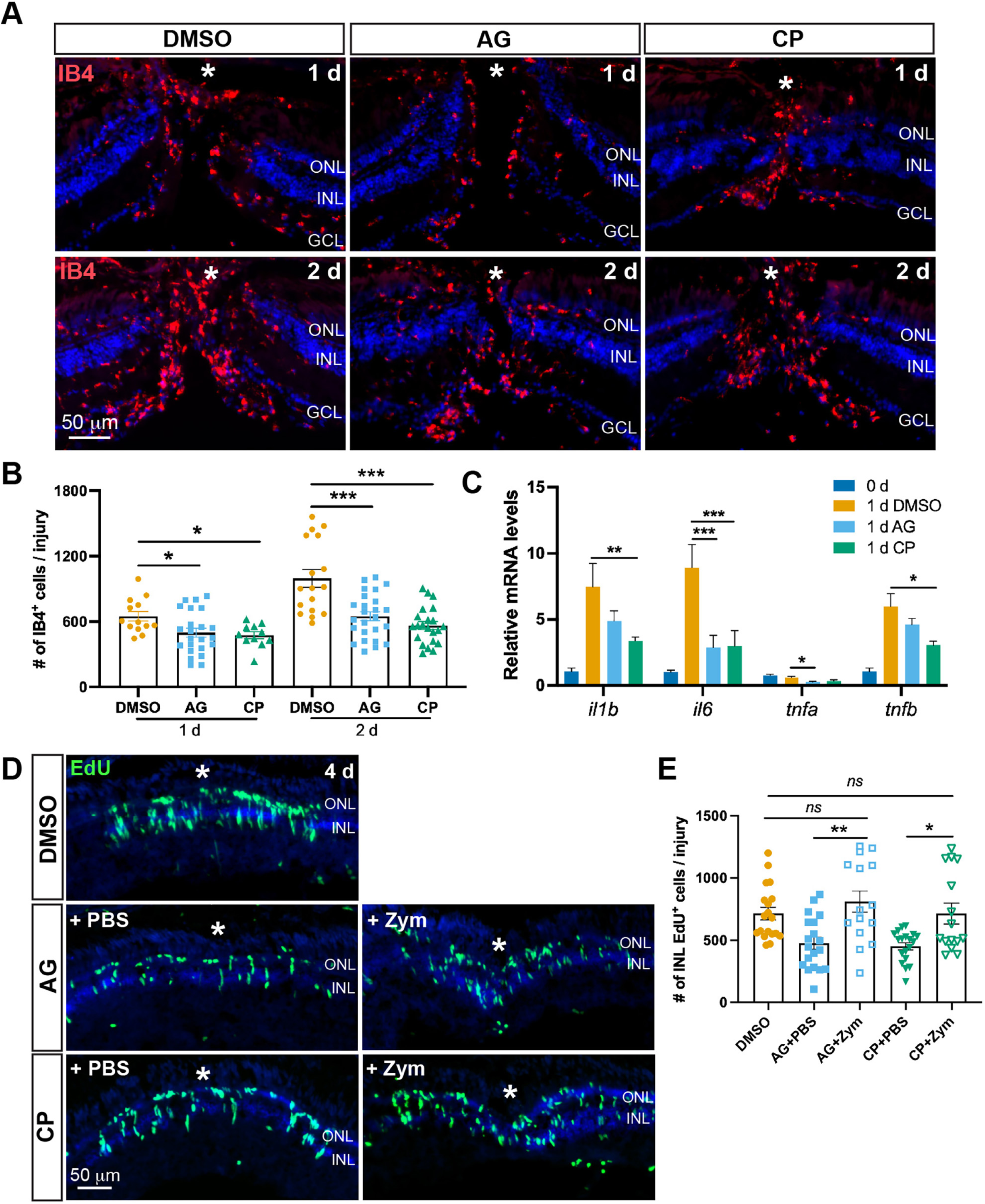
Neutrophil presence promotes microglia reactivity and the inflammatory response in stab-injured retinas. (A) Fluorescence microscopy showing the presence of IB4^+^ microglia in DMSO- or AG/CP-treated retina at 1 and 2 dpi. (B) Quantification of microglia number in (A). (C) qPCR analysis showing the mRNA expression of typical inflammatory cytokines in uninjured control or 1 dpi DMSO/AG/CP-treated retinas. (D) EdU immunofluorescence showing the presence of proliferating MGPCs in retinas of indicated groups at 4 dpi. (E) Quantification of the number of INL EdU^+^ cells per injury in (D). White asterisks, site of the needle-poke injury. ONL, outer nuclear layer; INL, inner nuclear layer.

### 3.7 Neutrophils may promote the retinal regenerative response through Hb-egfa expression

In addition to immune modulation, neutrophil could also regulate MG reprogramming by secreting regeneration-associated growth factors. Because *hbegfa* was the only growth factor gene expressed by neutrophils in our scRNA-seq dataset (Figure 1E), and a previous study has reported that MG-derived Hb-egfa plays a critical role in MG dedifferentiation and retinal regeneration(27), we further investigated the expression and function of Hb-egfa in stab-injured zebrafish retinas. scRNA-seq showed that *hbegfa* was mainly expressed by neutrophil, MG subtypes including activated MG (Act-MG) and proliferating MG (Pro-MG), and endothelial cells (EC), whereas microglia express the other Hb-egf homolog *hbegfb* (Figure 7A,B). scRNA-seq further demonstrated that Egfra (encoded by *egfra*) was the main Hb-egf receptor expressed in the retina, whereas other Hb-egf receptors such as Erbb4a and Erbb4b were barely detected (Figure 7A,C). Interestingly, *egfra* was weakly expressed in MG but enriched in EC, suggesting that Hb-egfa may regulate MG reprogramming in a non-cell-autonomous manner. Because neutrophil invasion is a very early event, we speculate that neutrophils might be the main source of retinal Hb-egfa during the early phase after injury, earlier than that expressed by activated MG or proliferative MG, which are typically detected at 1 or 2 dpi, respectively. Indeed, RT-PCR showed that *hbegfa* expression was induced in the retina as early as 1.5 hpi, earlier than the MG activation marker *ascl1a* (Figure 7D). In contrast to the continuous increase of *ascl1a* expression from 0 to 48 hpi, *hbegfa* expression peaked at around 3 to 6 hpi, decreased from 12 to 24 hpi, and rebounded at 48 hpi (Figure 7D). This expression pattern aligns with the neutrophil peak at the injury site at 3-6 hpi. Importantly, neutrophil depletion by AG or CP treatment drastically reduced retinal *hbegfa* levels at 3, 12 and 24 hpi as shown by qPCR analysis (Figure 7E), consistent with the idea that neutrophils are the main source of *hbegfa* during the early hours following retinal injury.

**Figure 7.**
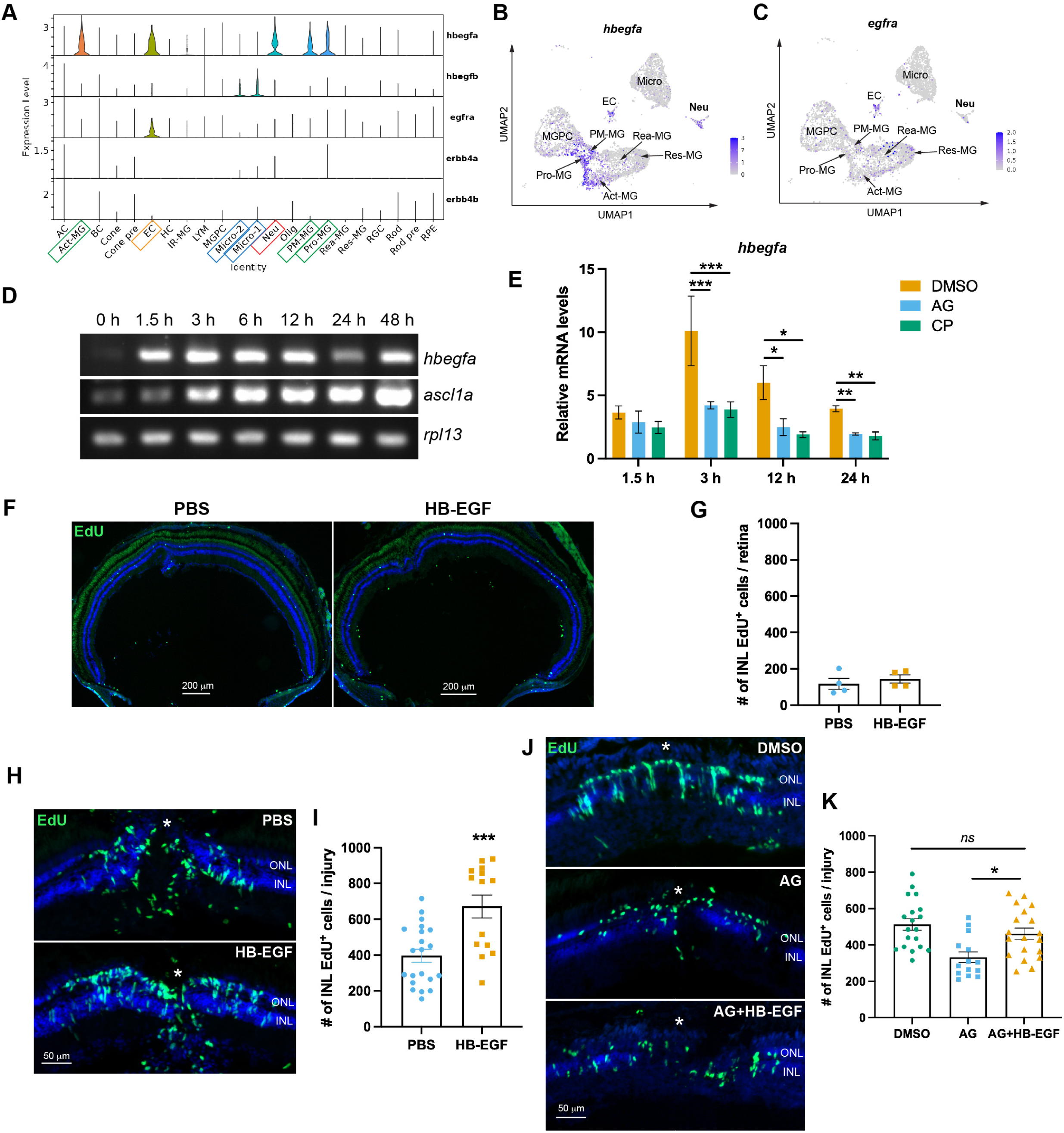
Neutrophils express a regeneration-promoting factor Hb-egfa in stab-injured retina. (A-C) scRNA-seq analysis showing the expression of *hbegfa*, *hbegfb* and their receptors (*egfra*, *erbb4a* and *erbb4b*) in each retinal cell types/subtypes. AC, amacrine cell; Act-MG, activated MG; BC, bipolar cell; Cone pre, cone precursor; EC, endothelial cell; HC, horizontal cell; IR-MG, immunoregulatory MG; LYM, lymphocyte; Micro, microglia; Neu, neutrophil; Olig, oligodendrocyte; PM-MG, post-mitotic MG; Pro-MG, proliferating MG; Rea-MG, reactive MG; Res-MG, resting MG; RGC, retinal ganglion cell; Rod pre, rod precursor. (D) RT-PCR showing the expression of *hbegfa* and *ascl1a* at indicated timepoints in stab-injured zebrafish retina. *rpl13* serves as the house-keeping control. (E) qPCR showing the expression of *hbegfa* at indicated timepoints in DMSO- or AG/CP-treated retinas. (F) EdU immunofluorescence assessing cell proliferation in PBS- or HB-EGF-injected uninjured retina at 2-day post injection. (G) Quantification of the number of INL EdU^+^ cells per retina in (F). (H) EdU immunofluorescence showing the presence of proliferating MGPCs in the PBS or HB-EGF-injected injured retina at 4 dpi. (I) Quantification of (H). (J) EdU immunofluorescence showing the presence of MGPCs in stab-injured retinas that received indicated treatments. (K) Quantification of (J). White asterisks, site of the needle-poke injury. ONL, outer nuclear layer; INL, inner nuclear layer.

We next examined the effect of HB-EGF over-expression on the regenerative response in uninjured or stab-injured zebrafish retinas. In uninjured retinas, daily intravitreal injection of a recombinant human HB-EGF protein (200 ng per eye) for 2 days did not stimulate MG proliferation (Figure 7F,G). In contrast, a single intravitreal HB-EGF injection (200 ng) at the time of injury significantly increased the number of MGPCs at 4 dpi in stab-injured retinas (Figure 7H,I). These results demonstrate that HB-EGF can promote the regenerative response in the injured retina. To investigate whether neutrophils regulate the regenerative response through this growth factor, neutrophil-depleted retinas were treated with HB-EGF (200 ng) for 2 days, and EdU immunofluorescence showed that HB-EGF injection restored MPGC numbers in AG-treated retinas to levels comparable to those in untreated controls (Figure 7J,K). Taken together, these findings suggest that neutrophils may regulate retinal regenerative responses through Hb-egfa expression.

## 4 DISCUSSION

In this study, we identify a previously unrecognized requirement for neutrophils in retinal regeneration and reveal a two-tiered control of their function in this process. The first tier is access: neutrophils are required for regenerative responses after stab injury, where barrier disruption permits their entry into the retinal parenchyma, but not after NMDA injury, where they remain excluded and their depletion has no effect. The second tier operates once access is granted: Cxcl12-Cxcr4b signaling retains neutrophils at the injury site, and manipulating this retention correspondingly alters MG proliferation, MGPC formation, and neuronal regeneration. Within this retention window, neutrophils modulate microglial reactivity and inflammatory responses and provide Hb-egf as a regeneration-promoting factor. These findings position neutrophils as context-dependent regulators of retinal regeneration and suggest that access and local retention, rather than cell identity alone, determine whether and how they contribute to repair.

The differential access of neutrophils to the injured retina is most simply explained by the barrier status of the two paradigms. White et al. previously showed that neutrophils infiltrate the retina after puncture injury but display no reactivity to selective rod cell loss in zebrafish larvae, and interpreted this difference in light of BRB disruption(15). Our ablation experiments extend this observation from a correlative description to a functional consequence: when neutrophils are permitted to enter after stab injury, their depletion impairs regeneration; when they are excluded in the NMDA model, depletion has no effect. This establishes that access is not merely a correlate of injury type but a variable with regenerative consequences. We note, however, that BRB permeability was not directly measured in our study, and injury-specific cues independent of barrier status may also contribute to the differential access we observe. Distinguishing these possibilities will require direct assessment of barrier integrity across paradigms, but the access framework provides a testable entry point for that question.

If access constitutes the first tier, the second tier concerns the behavior of neutrophils once they have entered. Our Cxcl12-Cxcr4b experiments address the retention dimension of this tier. Cxcl12-Cxcr4 signaling is known to retain neutrophils at inflammatory sites in zebrafish(37), and our data show that manipulating this axis shifts neutrophil numbers specifically during the resolution phase without affecting initial infiltration. Reducing retention with AMD3100 decreased MG proliferation and MGPC formation, whereas enhancing retention with Cxcl12a increased both. This positions Cxcl12-Cxcr4b signaling as a molecular handle on neutrophil persistence at the injury site, rather than a determinant of recruitment. Importantly, the relationship between retention and regeneration in our system appears monotonic within the range tested: greater retention yielded greater regenerative output. This contrasts with the spinal cord, where promoting neutrophil resolution through Cxcr4 inhibition enhances regeneration, and blocking recruitment through Cxcr1/2 inhibition impairs it(24). In that context, neutrophils are required, but their timely departure is also necessary; excessive persistence is detrimental. The difference between these outcomes likely reflects tissue-specific requirements for the duration of neutrophil activity, and it reinforces the view that retention is a tunable variable whose optimal set point is context-dependent rather than fixed.

Within the retention window defined by Cxcl12-Cxcr4b signaling, neutrophils appear to influence regeneration through at least two functional outputs. First, neutrophil depletion reduced microglial accumulation at the injury site and lowered expression of *il1b*, *il6*, *tnfa*, and *tnfb*, consistent with neutrophils acting upstream of microglial reactivity and the inflammatory response in the stab-injured retina. This aligns with observations in the regenerating spinal cord, where neutrophil-specific transcriptomic profiling correlates with transient tnf-α induction in macrophage/microglia populations, and where blocking neutrophil recruitment diminishes macrophage/microglia presence at the injury site(24). Together, these findings suggest that a neutrophil–microglial signaling axis may be a shared feature of regenerating zebrafish tissues, though the direction and molecular mediators of this crosstalk remain to be defined in the retina. Second, neutrophils provide Hb-egf. Our scRNA-seq data identified *hbegfa* as the only growth factor gene detected in retinal neutrophils, and neutrophil depletion drastically reduced retinal *hbegfa* levels at early time points. Exogenous HB-EGF restored MGPC formation in neutrophil-depleted retinas, supporting Hb-egf as a functionally relevant output. This places neutrophil-derived Hb-egf within the established HB-EGF/EGFR/MAPK/Ascl1a circuitry that governs MG dedifferentiation(27), but with a distinct cellular source: whereas MG themselves are a major source of Hb-egf at the injury site, neutrophils provide an earlier bolus that may help initiate the response before MG-derived Hb-egf sustains it. The same ligand has been implicated in neutrophil-dependent epicardial regeneration in the zebrafish heart(25), where *hbegfa* released from neutrophils synergizes with FGF/MAPK signaling, raising the possibility that neutrophil-derived Hb-egf is a recurrent regenerative signal across tissues.

An important boundary of our framework is that access and retention are not proposed as sufficient to explain all context-dependent outcomes of neutrophil function. Neutrophils have been reported to impede regeneration in other settings: in wound-induced hair follicle neogenesis, mature neutrophils and their neutrophil extracellular traps (NET) hinder regeneration, and their depletion enhances regenerative outcomes(38). In the regenerating fin, neutrophils are dispensable(39). These examples indicate that entry into a tissue does not guarantee a pro-regenerative contribution, and that additional state-dependent variables—such as the molecular profile of recruited neutrophils, the duration and intensity of NET formation, and the local cellular interlocutors—can shift the net effect toward inhibition. Our study identifies access and retention as two variables that are necessary to consider when interpreting whether neutrophils contribute positively to a given regenerative process; whether the same variables account for inhibitory outcomes, or whether distinct state-dependent programs predominate in those settings, remains an open question.

Several limitations should be acknowledged. First, our pharmacological ablation strategy using Erbb2 inhibitors, while effective in reducing neutrophil numbers, may have off-target effects on other Erbb2-expressing cells. However, treatment using the same Erbb2 inhibitors had no significant effect on the regenerative responses in the NMDA-injury model, suggesting that this is unlikely the case. Nevertheless, genetic ablation models or neutrophil-specific Cre or nitroreductase lines would strengthen the conclusions. Second, the precise cellular target of neutrophil-derived Hb-egf remains unresolved. scRNA-seq indicated that *egfra* is weakly expressed in MG but enriched in endothelial cells, raising the possibility that Hb-egf acts on the vasculature or on another intermediate cell type rather than directly on MG. Conditional manipulation of Egfra in specific retinal cell types will be required to resolve this. Third, we did not directly measure BRB permeability in the two paradigms, and the access interpretation rests on the observed exclusion of neutrophils from the NMDA-injured parenchyma rather than on a direct barrier assay. Fourth, whether the access–retention framework extends to other BRB-disrupting injuries, such as laser damage or vascular injury, and whether it can be harnessed to prime mammalian MG for reprogramming, are questions that follow directly from these findings but remain to be tested.

In summary, our work identifies neutrophils as context-dependent regulators of retinal regeneration and organizes their contribution into two experimentally dissociable tiers: access to the injured tissue and local retention that enables regenerative signaling. This framework accommodates the divergent roles of neutrophils across regenerative and non-regenerative settings, and it suggests that manipulating access or retention, rather than targeting neutrophils indiscriminately, may offer a more precise strategy for modulating immune contributions to CNS repair.

## ACKNOWLEDGEMENT

This study was supported by grants from NSFC (81970820), and Basic Research Program of Jiangsu Education Department (24KJA180008).

